# Tempo-dependent selective enhancement of neural responses at the beat frequency can be mimicked by both an oscillator and an evoked model

**DOI:** 10.1101/2024.07.11.603023

**Authors:** Atser Damsma, Mitchell de Roo, Keith Doelling, Pierre-Louis Bazin, Fleur L. Bouwer

## Abstract

A crucial mechanism for the brain to make sense of the auditory environment is the synchronization of neural responses to external temporal regularities, such as a musical beat. It is debated whether this synchronization and the resulting beat percept reflect phase alignment of endogenous neural oscillations to the external regularity (*entrainment*), or evoked responses to the rhythmic stimulus (*tracking*). Here, we use the tempo-dependent properties of beat processing to differentiate between the two accounts. Participants listened to a repeating rhythmic pattern at different speeds. Behaviorally, they consistently tapped at the preferred beat rate (around 2 Hz) across tempi, shifting to higher metrical levels as tempo increased. We found a similar shift in EEG data, where the metrical level at which neural synchronization was strongest depended on tempo. This selective enhancement is consistent with entrainment accounts and could indeed be mimicked by an oscillator model. However, importantly, the results were also captured by a model simulating evoked responses. Together, our findings demonstrate that while neural responses to rhythm are selectively enhanced at the beat rate, this enhancement need not be taken as evidence for entrainment, but can also be explained by successive evoked responses.

---

Picking up on temporal structure in music is crucial for the anticipation of upcoming sounds, and for synchronizing movements between individuals (Keller et al. 2014; Bouwer and Honing 2015). The extraction of a regular beat – a subjective periodic pulse perceived in a sound pattern – is fundamental to this process (Large and Palmer 2002; Honing 2012). In addition, a hierarchy of regularly alternating stronger and weaker beats can be perceived, known as *meter*. Beat and meter perception are often interpreted in the light of Dynamic Attending Theory (DAT), which proposes that fluctuations in attention synchronize to a regular stimulus in the environment, creating peaks and dips in processing sensitivity that coincide with the regularity of a beat (Large and Jones 1999; Large and Palmer 2002). This synchronization, or *entrainment*, has been proposed to rely on coupling of low-frequency neural oscillations to an external stimulus (Schroeder and Lakatos 2009; Henry and Herrmann 2014). Oscillatory entrainment of neural populations could indeed be an efficient implementation of temporal expectations in the brain, aligning phases of optimal sensitivity to the timing of upcoming stimuli (Schroeder and Lakatos 2009; Henry and Obleser 2012; Arnal et al. 2015; Zoefel et al. 2018). Although there is evidence that neural dynamics synchronize to rhythmic stimuli, it is unclear whether this evidence reflects actual oscillatory mechanisms, or rather, passive neural tracking of incoming sounds (Obleser and Kayser 2019).

Brain responses related to beat and meter have been shown to be enhanced compared to the frequency spectrum of the rhythmic stimuli themselves (Nozaradan et al. 2012; Nozaradan 2014), or to a cochlear model of early auditory processing (Lenc et al. 2018; Lenc et al. 2022; Lenc et al. 2023). This selective enhancement of metrical frequencies has been especially apparent for rhythms that are only weakly metrical, or that do not even contain these frequencies themselves (so-called “missing-pulse” rhythms; Large et al. 2015; Tal et al. 2017), potentially reflecting a ‘periodization’ of the input towards a metrical representation in the brain (Nozaradan et al. 2012; Lenc et al. 2018; Nozaradan, Keller, et al. 2018; Lenc et al. 2021). Although it has been posited that this mechanism is unlikely to result from low-level processing (Large et al. 2015; Nozaradan, Schönwiesner, et al. 2018; Lenc et al. 2021), interpreting selective enhancement of beat-related frequencies as an oscillatory internal representation of the beat is not straightforward (see also Henry et al. 2017). Since every sound can be assumed to elicit an auditory evoked response in the EEG signal, any rhythmic stimulus could be expected to result in a correspondingly rhythmic cortical response, which may be difficult to distinguish from endogenous oscillatory activity (Zoefel et al. 2018; Doelling et al. 2019; Obleser and Kayser 2019). Indeed, it has been argued that neural entrainment measured in response to rhythmically changing visual and speech stimuli can likely be explained by mere transient event-related potentials (Capilla et al. 2011; Novembre and Iannetti 2018; Oganian et al. 2023), although these linear responses do not readily explain the missing pulse phenomenon. Thus, the question whether oscillatory entrainment indeed underlies the extraction of a regular beat from a time-varying rhythmic signal is still open.

One way to disentangle oscillatory entrainment from evoked responses is to compare brain responses to stimuli presented at different rates (Capilla et al. 2011; Doelling et al. 2019; Oganian et al. 2023). The beat is considered the most salient regularity that is perceived in a rhythm (Honing and Bouwer 2019), and within nested levels of regularity often present in rhythm (the metrical structure), the hierarchical level at which a beat is perceived depends on the tempo at which a rhythm is played (Parncutt 1994; Drake et al. 2000). Humans typically show a preference for beat rates between 1.25 and 2.5 Hz (Fraisse 1982; van Noorden and Moelants 1999; Moelants 2002; Zalta et al. 2020; Weineck et al. 2022), corresponding to common tempi in popular music (Moelants 2002) and spontaneous body movements (MacDougall and Moore 2005; McAuley et al. 2006). When confronted with rhythms at faster rates, attention shifts to regularities at a slower rate (i.e., a higher level in the metrical hierarchy) (Parncutt 1994; Drake et al. 2000). If cortical responses represent entrainment to the perceived beat, we may therefore expect tempo-dependent selective enhancement of beat frequencies, while they may be tempo-invariant if they directly follow the sound signal.

In addition, oscillatory and evoked accounts of responses to rhythm make different predictions regarding the relationship between the phase of the EEG signal and the phase of the sound signal. For example, Doelling et al. (2019) presented participants with clips of piano music at six different note rates, while measuring magnetoencephalography (MEG). Manipulating speed in this way allowed them to make opposing predictions for the two hypotheses: while the fixed latency of evoked responses should lead to a phase shift in the EEG signal relative to sound onset over different stimulus rates, oscillatory activity is hypothesized to adapt to the stimulus rate and therefore result in a relatively stable phase relationship. Indeed, Doelling et al. showed that an oscillator model predicted larger phase stability, defined in a phase concentration metric (PCM), compared to an evoked model. The PCM of the MEG signal was shown to be higher than could be predicted by evoked responses alone, leading the authors to conclude that the cortical signal contained contributions from oscillatory entrainment. However, this study focused not on beat perception but on temporal prediction of individual notes based on a unimodal distribution of temporal intervals centered on a specific note rate. As such, natural music clips were selected for isochronicity, which induce both beat perception as well as temporal expectations based on the repeated single interval (Bouwer et al. 2021; Bouwer et al. 2023). Furthermore, as a result of the project’s aims, entrainment was considered only at the note rate, which was in some cases much faster than the beat rate that humans prefer. Hence, it is unclear whether these findings would also apply to hierarchical beat processing (Honing 2012).

In addition to the question of whether cortical responses to rhythm constitute true oscillatory entrainment, it is yet unknown whether neural entrainment is an automatic mechanism or instead depends on top-down control (Rimmele et al. 2018; Obleser and Kayser 2019; Bouwer 2022). While temporal prediction and beat perception have been shown to occur under passive listening conditions (Bouwer et al. 2014; Damsma and van Rijn 2017; Bouwer et al. 2020) oscillatory neural entrainment may depend on attention. For example, in monkeys, entrainment in the auditory cortex was found only for attended rhythmic auditory streams, but not when the stimulus was ignored (Lakatos et al. 2013). In humans, sustained neural oscillations have been demonstrated after the offset of task-relevant rhythmic speech (Kösem et al. 2018; van Bree et al. 2021; Bouwer et al. 2023), whereas passive listening conditions did not yield similar ongoing entrainment in the delta range (Lerousseau et al. 2021), suggesting that it might be a task-dependent phenomenon under top-down control (Bouwer 2022).

The current study aimed to investigate whether cortical responses to time-varying rhythm show evidence of entrainment by testing how they change over tempo and task demands. Participants were presented with a simple non-isochronous rhythm played at different rates, while EEG was measured in two conditions: a passive and an active listening condition. In addition, participants were asked to tap along to the beat in these rhythms on separate trials. Behaviorally, we expected that the most salient level of regularity (i.e., the beat) would shift depending on the tempo, to retain a perceived beat close to preferred rate (i.e., ∼2 Hz). Thus, we expected participants to tap at the grid rate (i.e., the rhythmic pattern could be placed on a duple-meter grid) at slower tempi, but at half the grid rate at faster tempi. To test whether the tempo-dependency of behavioral markers of beat perception is supported by oscillatory entrainment, first, we investigated whether the spectral power and phase coherence of the EEG signal showed similar tempo-dependent shifts to the preferred beat range. Such shifts would be apparent from selective enhancement of the EEG signal at the beat rate. Second, brain responses were simulated using two competing models: an oscillator model and a model of evoked responses. We expected the output of the oscillator model, but not the evoked model, to show selective enhancement at the beat frequency, and thus, to better capture the EEG data. Third, phase concentration of the EEG data was compared to the predictions of these two models. As in previous work (Doelling et al. 2019), we expected the models to make diverging predictions, with the output of the oscillator model having a higher phase stability (i.e., a higher PCM) than the output of the evoked model. We expected the EEG data to have a higher phase stability than could be expected from the evoked model alone. Finally, we tested whether the power and phase of neural entrainment depend on attention, by contrasting active and passive listening conditions.

## Materials and Methods

### Experimental design

#### Participants

Twelve participants (5 women; age range, 21-31 years, M = 27, SD = 4.6, age of one participant unknown) took part in the EEG experiment after providing written consent and were rewarded a monetary compensation (€22.50). None of the participants reported hearing problems or a history of neurological disorders. The experiment was approved by the ethics review board of the Faculty of Humanities of the University of Amsterdam.

#### Stimuli

The auditory stimuli consisted of woodblock sound generated using GarageBand (Apple inc.) arranged in a simple, non-isochronous rhythmic pattern in duple meter (Figure 1). The rhythmic pattern was designed so that the duration between grid points was equal (the *grid interval*), and each grid point contained a sound or a silence. The pattern always consisted of a sound, a silence, and two sounded grid points (see Figure 1). Five different tempi were presented using grid intervals of 150, 200, 300, 400, and 600 ms. Given that the preferred beat rate is close to ∼2 Hz, the perceived beat was expected to shift from occurring at every complete pattern of four grid points for fast tempi (e.g., the 150 ms grid) to occurring at every two grid points or at every single grid point for slower tempi (e.g., for the 300 ms and 600 ms grid, respectively). Note that the expected perceived regularities at the 150, 300, and 600 ms tempi were nested, as were the expected perceived regularities at the 200 and 400 ms tempi.

**Figure 1.**
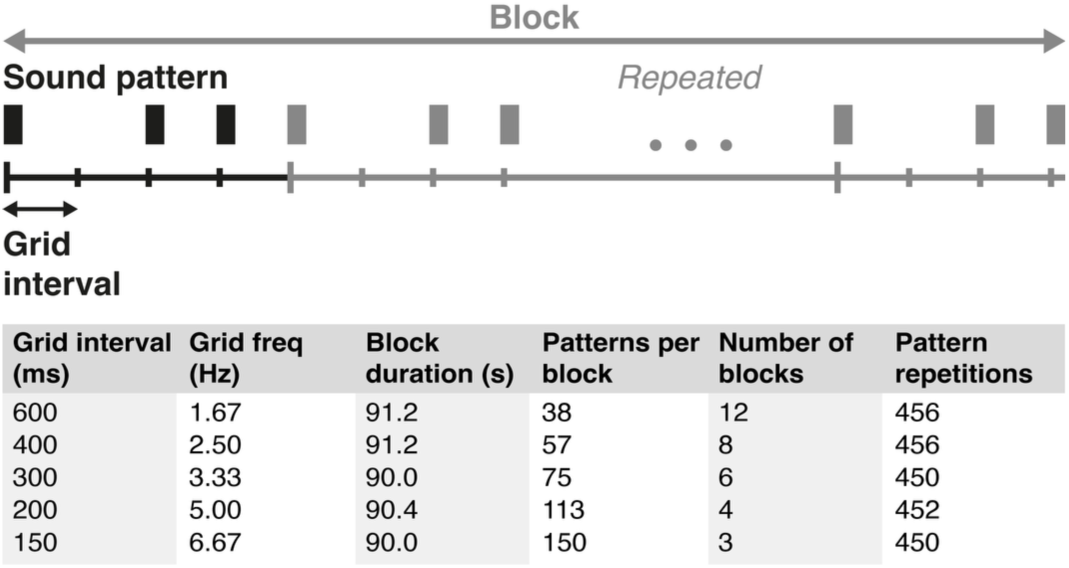
Overview of the auditory stimuli and experiment design. A woodblock sound (depicted here as a black rectangle) was presented in a simple, non-isochronous rhythmic pattern that could be placed on an imaginary duple-meter grid. The pattern was presented in five different tempi over separate blocks in the experiment, with corresponding grid intervals ranging from 150 to 600 ms. In a block, the rhythmic pattern was repeated several times. The number of repetitions was dependent on the tempo, but always resulted in a minimum block duration of 90 s. The blocks were repeated several times over the course of the experiment, so that the total number of pattern presentations was as equal as possible between the five different tempo conditions. The number of blocks and total pattern repetitions presented here are for each of the two attention conditions (i.e., the total number of pattern repetitions was >900 for each tempo).

#### Procedure

The rhythmic pattern was repeated within blocks to form continuous sequences. The number of repetitions per block and the number of blocks were adjusted to the grid interval, so that the minimum duration of each block was 90 s and each rhythmic pattern was presented a minimum of 450 times in each attention condition (Figure 1). This resulted in a minimum of 900 total pattern repetitions per tempo over the two attention conditions in the experiment. Auditory playback of the pattern introduced a small systematic delay between repetitions (range: 4.6-7.1 ms), which was corrected for in subsequent analyses. Each block started with a 5 s baseline period before sound onset. Block order was random for each participant. Participants were allowed to take short breaks in between blocks.

Participants were presented with two different conditions: an *unattended* and an *attended* condition. Participants always started with the unattended condition, and were allowed a break before the start of the attended condition. In the unattended condition, participants watched a silent movie with subtitles, while instructed to ignore the rhythms and keep attention to the movie. Participants were free to select a movie from a selection of available DVDs. In the attended condition, there was no movie playing and participants were asked to pay attention to the rhythms. In between randomly selected blocks in the attended condition, for a total of 10 times, participants received one of the following two questions: “How fast do you think the last rhythm you heard sounded?” and “How easy do you think it would be to dance to the last rhythm you heard?”. Participants were asked to answer the question on a scale from 1-10. The questions were only included to engage the participants with the rhythms and are not analyzed here. After the experiment, participants were asked to fill in a questionnaire on their educational and musical background.

### Tapping

#### Tapping recording

At the end of the experiment, participants tapped along with the repeating pattern in a separate trial for each tempo. Each trial was a minimum of 25 seconds (the precise number of pattern repetitions depended on the grid frequency). Taps were recorded using a button box. Participants were instructed to tap along regularly, the way they would tap their foot to the rhythm. If participants synchronized to the pattern instead of the beat, the trial was discarded, and they were explained what the beat is. Then, they could try tapping with their foot first, before resuming tapping with their index finger to record the data.

#### Tapping analysis

During pre-processing, recorded taps that followed quickly (i.e., within 50 ms) after another tap were removed, as these likely constituted technical noise instead of valid taps. To make the analysis comparable to the EEG data, a 512 Hz time series of zeros was created in which tap onsets were coded as 1, starting with the first tap at time point 0. Per participant and tempo condition, spectral power was calculated using a Fast Fourier transform (FFT). For the FFT, each time series was zero-padded or truncated to 24 s, corresponding with a frequency resolution of 0.042 Hz. This resolution was chosen to (1) extract power at all frequencies of interest, (2) include all taps (with one single exception in one participant), and (3) make the spectral output comparable across the tapping, EEG, and model data. Next, the power at every frequency point was averaged with its two neighboring frequencies to account for imprecision in frequency-specificity. Meter-related and meter-unrelated frequencies were marked as frequencies of interest. For each tempo, meter-related frequencies were defined as the grid frequency, first subharmonic (i.e., grid frequency/2), and second subharmonic (i.e., grid frequency/4). Note that for the faster tempi, this meant that some of the meter-related frequencies were identical to the grid frequencies of the slower tempi (e.g., the first subharmonic of the 150 ms grid was identical to the grid frequency of the 300 ms grid). Meter-unrelated frequencies were in turn defined as the meter-related frequencies of unrelated tempo conditions (i.e., 200 and 400 ms for the 150, 300, and 600 ms conditions, and vice versa). Next, for each participant and tempo condition, the power at frequencies of interest was *z*-scored. Specifically, we computed the mean and standard deviation of power over all frequencies of interest. For each frequency of interest, we then calculated *z*-scored power by subtracting the mean and dividing by the standard deviation. Finally, we averaged the *z*-scored power over the different meter-unrelated frequencies.

A linear mixed-effects model (LMM) with *power* (i.e., the *z*-scored power values) as the dependent variable was computed using the *lme4* package in *R* (Bates et al. 2015). *Tempo* (600 ms, 400 ms, 300 ms, 200 ms, or 150 ms), *frequency* (grid frequency, first subharmonic, second subharmonic, meter-unrelated), and their interactions were entered as categorical fixed factors and *participant* as a random intercept. Omnibus effects of the fixed factors were estimated with type III Wald Chi-Square tests using the *Anova* function from the *car* package in *R* (Fox et al. 2012). Contrasts between conditions were assessed using Tukey’s pairwise multiple comparison test with corresponding effect size analysis using the *emmeans* package in *R* (Lenth 2023).

### EEG

#### EEG recording

EEG data was recorded at 512 Hz using a 64-channel Biosemi Active-Two acquisition system (Biosemi, Amsterdam, The Netherlands) according to the to the standard 10/20 configuration. Additionally, electrodes were recorded from EOG channels, mastoids and the nose.

#### EEG pre-processing

EEG pre-processing was performed using EEGlab (Delorme and Makeig 2004) and Fieldtrip (Oostenveld et al. 2011) in Matlab R2021a (Mathworks). EEG data was re-referenced to the average. The data was manually checked for bad channels, which were replaced by data interpolated from surrounding channels. The average number of interpolated channels per participant was 1.17. However, none of these channels were used in the eventual analysis. Eye blinks were removed using independent component analysis (ICA). Finally, a 0.01 Hz high-pass filter was applied to remove slow drift.

### EEG data analysis

#### FFT on EEG data

First, we calculated spectral power in the EEG signal by performing an FFT to assess whether frequencies at the beat level were selectively enhanced. To improve the signal-to-noise ratio, shorter epochs of 12 s were created for all tempo conditions. The epochs always started at pattern onset and were non-overlapping. The resulting epochs were averaged per participant, attention condition, and tempo, similar to the procedure followed in previous research to assess phase-locked activity (Nozaradan et al. 2011; Nozaradan et al. 2012; Nozaradan et al. 2015). Next, per electrode, the data was demeaned and the FFT was calculated with the data zero-padded to a length of 24 s to match the 0.042 Hz frequency resolution of the tapping and model analysis. Finally, the power spectrum was calculated as the average EEG signal at a fronto-central electrode cluster (Fz, F3, F4, F1, F2, Fcz, Fc3, Fc4, Fc1, Fc2, Cz, C3, C4, C1, C2), which has been shown to reflect maximum auditory synchronization (Nozaradan et al. 2011; Nozaradan et al. 2015). Task-unrelated background activity was removed by subtracting, from each point in the power spectrum, the average power at two frequency points surrounding, but not directly neighboring, the frequency of interest (−2 and +2, corresponding to steps of −0.083 Hz and +0.083 Hz) (Nozaradan et al. 2011; Nozaradan et al. 2012; Nozaradan et al. 2015). This procedure is based on the assumption that neural synchronization is highly frequency specific, while background noise is similar across neighboring frequencies.

Statistical analysis of the frequency-domain EEG was similar to the tapping data analysis: first, the power at every frequency point was averaged with its two immediately neighboring frequencies (−1 and +1) to account for imprecision in the frequency-specificity of the synchronization. Then, the relative power at meter-related and meter-unrelated frequencies was calculated per participant, attention condition, and tempo. An LMM with this *z*-scored *power* as the dependent variable was performed with *tempo* (600 ms, 400 ms, 300 ms, 200 ms, or 150 ms), *attention* (attended or unattended), *frequency* (grid frequency, first subharmonic, second subharmonic, meter-unrelated), and their interactions as categorical fixed factors and *participant* as random intercept. Omnibus effects and contrasts were estimated in the same way as for the tapping data.

If neural synchronization underlies beat perception, we may expect that participants’ EEG power predicted their preferred tapping rate. To test this, we computed a linear regression model predicting relative tapping power, using relative EEG power as a predictor (in addition to *tempo*, *frequency*, and their interaction). The results can be found in Supplementary Materials section 1.

#### ITC of EEG data

As an additional analysis to assess the possible selective enhancement of the beat frequency, we examined inter-trial phase coherence (ITC), which is often used to quantify neural tracking of rhythmic signals (Doelling and Poeppel 2015; Cameron et al. 2019). ITC measures the uniformity of phase angles across epochs: when neural responses are more consistently synchronized to a rhythmic signal, this phase uniformity increases. In order to calculate ITC, we first performed a time-frequency analysis on EEG data of the full block. The time-frequency analysis was performed on all time points using a Morlet wavelet with a width of 7 cycles on frequencies of interest between 0.33 and 7.5 Hz (0.33, 0.42, 0.52, 0.63, 0.73, 0.83, 1.04, 1.25, 1.46, 1.67, 2.08, 2.50, 2.92, 3.33, 4.17, 5.0, 5.83, 6.67, and 7.50 Hz). Next, epochs in the time-frequency spectrum were defined corresponding to each repetition of the non-isochronous pattern (Figure 1). ITC was calculated as:

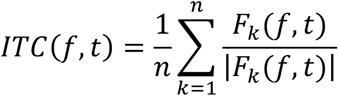

where *F*_k_(*f*,*t*) represents the complex spectral estimate of epoch *k* at frequency *f* and time *t*, and *n* is the number of epochs (Delorme and Makeig 2004). The possible range of the resulting ITC values is between 0 (high variability of phase angles across trials) and 1 (identical phase across all epochs). Finally, the ITC values were averaged over the cluster of electrodes of interest defined above for the FFT analysis.

Statistical analysis of the ITC was similar to the FFT analysis. ITC values were averaged over time, and then *z*-scored per participant, attention condition, and tempo. Tempo-related frequencies were defined as the grid frequency, first subharmonic, and second subharmonic. Frequencies which were not a harmonic of the grid frequency were labeled as meter-unrelated. An LMM was computed with relative *ITC* as dependent variable, with fixed and random factors, and omnibus and simple effects tests identical to the FFT analysis.

### Model simulations

To understand better how the selective enhancement observed in the EEG signal at the beat frequency arose, we used a modelling approach. We modelled the predicted EEG signal contrasting two models: an oscillator model and a model of evoked neural activity (Doelling et al. 2019).

#### Oscillator model

The oscillator model was based on the Wilson–Cowan model (Wilson and Cowan 1972; Doelling et al. 2019) (Figure 4A). The envelope of the continuous sound signals in every tempo condition served as input to the model. First, the envelope of the woodblock sound stimulus was obtained using a Hilbert transform (Matlab function *hilbert*), which was down-sampled to 512 Hz. For every tempo condition, the envelope of a full block of repeating patterns was created by convolving the woodblock envelope with a binary time series coding onsets. The resulting envelopes were normalized to serve as input to the model. The implementation of the model and its parameters was identical to Doelling et al. (2019), apart from a higher coupling value to accommodate a gain of the transient auditory signal (κ = 10) and a lower time constant (τ = 6.25), corresponding to a resting state frequency of ∼1.33 Hz. The resting state frequency was chosen to match the generally preferred beat period and the natural rate of spontaneous tapping (Zalta et al. 2020).

#### Evoked model

For the evoked model, a response kernel was computed using the Multivariate Temporal Response Function (mTRF) toolbox in Matlab (Crosse et al. 2016). This method derives a filter that maps the stimulus (i.e., the sound sequence) to the multi-channel EEG signal, where ridge regression is implemented to prevent overfitting. The ridge parameter was optimized using cross-validation, resulting in ***λ*** = 1. Cross-validation was performed on the full data set except for the 150 ms blocks, to prevent overfitting on the intervals shared between the 150, 300, and 600 ms conditions relative to the 200 and 400 ms conditions.

To determine the response kernel, a binary time series with impulses at sound onsets served as input to the mTRF algorithm. The EEG data was band-pass filtered between 0.1 and 30 Hz to exclude slow drifts and high-frequency noise. For each subject and attention condition, a response kernel was computed over the complete session. In this way, the kernel was forced to generalize over all tempo conditions and should not contain tempo-specific activity (such as tempo-specific oscillations). The time lag of the response filter was set to 0-1000 ms after sound onset. The response function was then averaged over the fronto-central electrode cluster that was also used for EEG analysis (Fz, F3, F4, F1, F2, Fcz, Fc3, Fc4, Fc1, Fc2, Cz, C3, C4, C1, C2), attention conditions, and participants. To prevent sudden jumps in the evoked model, the average response at time point 0 was used as a baseline and a half-Hann window was applied to the second half of the response kernel. For every tempo condition, an evoked model was then computed by convolving a binary time series with impulses at sound onsets with the response kernel (Figure 4D).

One possible pitfall of computing the response kernel over the entire session is that potential oscillatory activity may also be captured in the response kernels of corresponding tempo conditions (e.g., the evoked responses in the 300 ms condition may additionally contain overlapping oscillatory activity at 1.67 Hz by virtue of the presence of a beat). To confirm that the results were not dependent on such tempo-specific activity in the response kernel, additional response kernels were computed separately for frequency-unrelated tempo conditions (i.e., a response kernel estimated over data in the 150, 300, and 600 ms conditions was used for the evoked model of the 200 and 400 ms conditions, and vice versa). The results showed that an evoked model based on these kernels performed qualitatively similar to the model with the generalized response function described above, confirming that enhancements in the power spectrum are not due to frequency-specific oscillations in the response kernel. These results are reported in the Supplementary Materials section 2.

#### Comparing EEG to models

Subsequently, we compared the modelled outcomes with the actual EEG signal in two ways. First, we performed an FFT on the model output, similarly to the FFT on the EEG signal and the tapping data. Second, we computed the phase concentration for both the EEG data and the model outcomes.

#### FFT

A Fast Fourier transform was computed on the model output in a window of 2.4-26.4 s relative to the start of the block, so that block onset effects were reduced and the frequency resolution matched the spectral output of the EEG and tapping data. The FFT on the model output and subsequent normalization was similar to the tapping data. To quantify how well the models predicted the data, we calculated the mean squared error (MSE) between every participant’s *z*-scored EEG power at meter-unrelated frequencies, the grid frequency, first subharmonic, and second subharmonic and the power at these frequencies predicted by the oscillator model and evoked model. Because there was no evidence that neural synchronization was affected by attention, this calculation was performed on the average over attention conditions. Next, the MSE of the oscillator model and evoked model were compared using a paired samples Bayesian *t*-test using the *BayesFactor* package in *R* (Morey et al. 2014). The effect size (Cohen’s *d*) was estimated using the *rstatix* package in R (Kassambara 2023).

#### Phase concentration

The phase concentration metric (PCM, see Doelling et al. 2019) of the EEG signal was calculated based on the same 12 s epochs (averaged per tempo for each participant) used for the FFT analysis. Matching 12 s epochs were taken from the evoked model and oscillator model, starting from the third pattern repetition in a block to reduce onset effects. To obtain the PCM, all signals were filtered using a Gaussian filter in the frequency domain. For each tempo separately, the Gaussian distribution was centered at the grid frequency (e.g., 2.5 Hz in the 400 ms condition) with a standard deviation of 30% of this frequency (e.g., 0.75 Hz in the 400 ms condition). Next, phase at all pattern onsets during the epochs was obtained using a Hilbert transform (for an illustration, see Figure 4I). Note that this approach differs from Doelling et al. (2019), who employed a continuous phase measure: In our non-isochronous stimuli, pattern onset always aligned with the beat. To quantify phase difference, phase values of the stimulus envelope were subtracted from the phase values of the EEG, and the evoked and oscillator models (Doelling et al. 2019). The EEG phase differences were averaged per tempo condition and participant. Similarly, the average phase difference per tempo condition was calculated for the two models. The resulting phase vectors were normalized and averaged, and the PCM was calculated as the length of the average vector.

PCM values from the EEG signal were compared to the PCM of the two models using two separate Bayesian *t*-tests. Additionally, to test whether the participants’ PCM values were closer to one of the models, their squared error relative to the PCM of the models were compared using a paired samples Bayesian *t*-test. The primary phase concentration analysis was performed on the average EEG signal over attention conditions, as well as separately for the two conditions to test the effect of attention.

## Results

### Tapping results

Figure 2B shows the power spectrum of participants’ tapping, in which the grid frequency, first subharmonic (i.e., grid frequency/2), and second subharmonic (i.e., grid frequency/4) are highlighted in blue, red, and yellow, respectively. Figure S3A shows a histogram of the inter-tap intervals for each tempo and Figure S3B shows the relative power at meter-related and meter-unrelated frequencies for each participant. Participants predominantly tapped at the grid frequency for the slowest condition, but shifted their responses to higher metrical levels for faster tempi. Supporting this observation, the LMM predicting relative spectral power showed a significant *tempo x frequency* interaction (*χ*^2^(12) = 145.43, *p* < .001).

**Figure 2.**
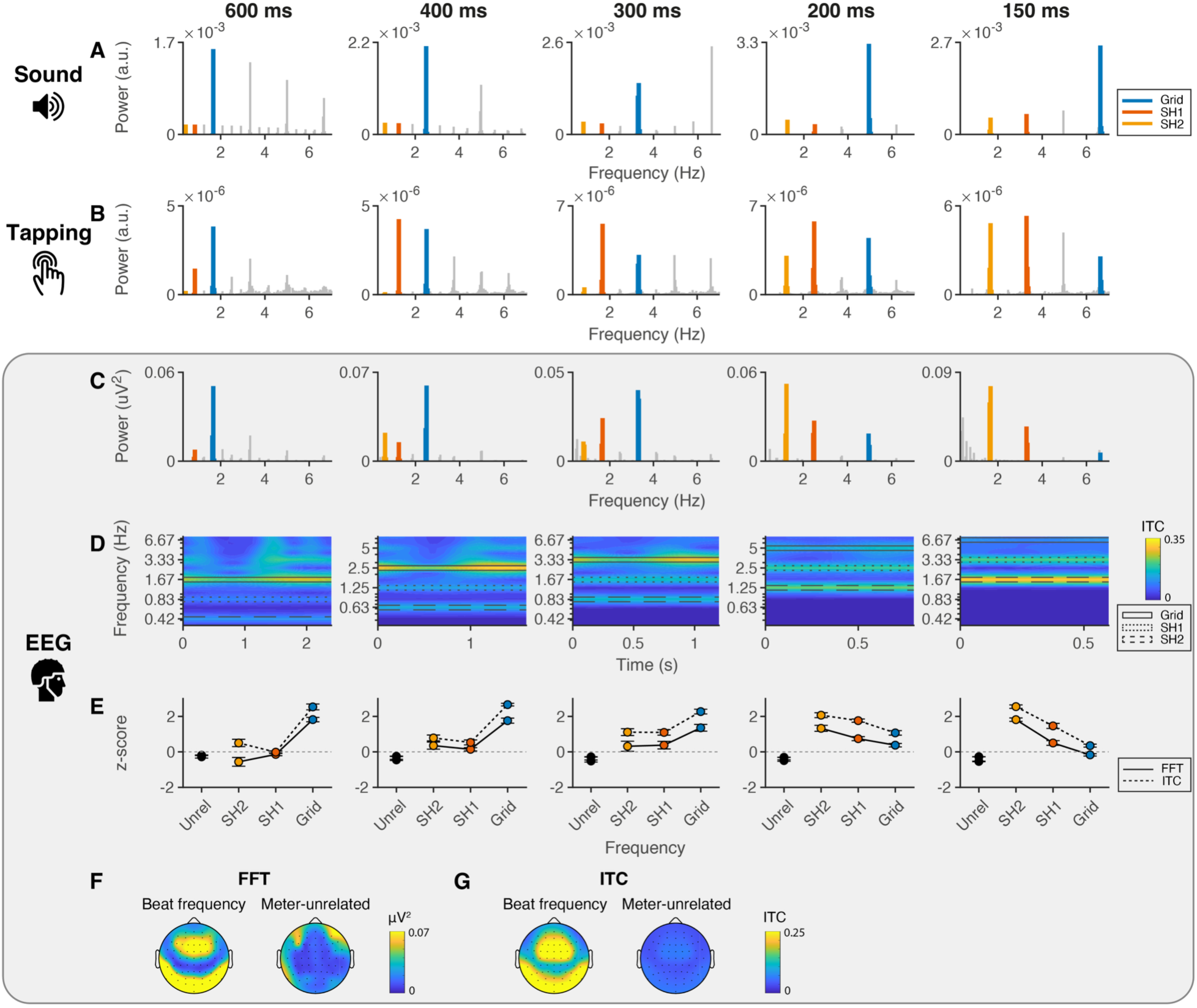
Spectral power shows tempo-dependent selective enhancement at the beat frequency. Spectral power of the sound, tapping and EEG signals over the five tempi, averaged across attention conditions. (A) Spectral power of the sound stimuli. Blue, red, and yellow colors highlight the grid frequency, first subharmonic, and second subharmonic, respectively. Note that, for visualization, the linewidth of meter-related frequencies has been increased and the y-axis range is adapted for each tempo. (B) Spectral power of the binary tapping signal over time. (C) Spectral power of the EEG signal after subtracting surrounding background noise. (D) Average inter-trial phase coherence (ITC) of the EEG signal. Rectangular boxes indicate meter-related frequencies. (E) Relative (z-scored) spectral power (FFT) and phase coherence (ITC) for meter-unrelated frequencies, the second and first subharmonics, and the grid frequency. Error bars represent standard error of the mean. (F) Topography of EEG power at the dominant beat frequency (left) and meter-unrelated frequencies (right). To display the average power at the dominant beat frequency, the frequency with maximum power was selected separately for each tempo. (G) Topography of ITC for the dominant beat frequency (left) and meter-unrelated frequencies (right).

Table S1 shows the full results of the post-hoc simple effects analysis with corresponding effect sizes. Power at the grid frequency was higher for the slowest (600 ms) condition compared to all faster tempo conditions (*p*s < .033), except the closest 400 ms tempo (*p* = .997). Similarly, it was lower for the 150 ms tempo compared to the 300, 400, and 600 ms conditions (*p*s < .041). Power at the first subharmonic, in contrast, was lower for the 600 ms tempo compared to the 200, 300, and 400 ms conditions (*p*s < .001). At the second subharmonic, relative power was higher for both the 150 and 200 ms conditions compared to the slower 300, 400, and 600 ms conditions (*p*s < .001). All other contrasts were non-significant (*p*s > .051).

In summary, as expected, these tapping results show a shift from synchronization at the grid frequency to lower subharmonics when the tempo was increased, which is consistent with the idea that listeners shift the metrical level of the perceived beat to stay close to their preferred rate.

### EEG results

#### FFT

Figure 2C shows the average spectral power at the fronto-central electrode cluster (Fz, F3, F4, F1, F2, Fcz, Fc3, Fc4, Fc1, Fc2, Cz, C3, C4, C1, C2) for each of the tempo conditions, Figure 2E shows the *z*-scored power at meter-related and meter-unrelated frequencies (note that the results depicted in Figure 2 are averaged over attention conditions), and Figure S4A shows the *z*-scored power per participant. Figure 2F shows the corresponding average EEG topography for the most prominent beat frequency and meter-unrelated frequencies. Similar to the tapping results, the power spectrum showed a shift from the grid frequency to lower subharmonics over increasing tempi. The LMM with relative power as the dependent variable indeed yielded a significant *tempo x frequency* interaction (*χ*^2^(12) = 235.35, *p* < .001), indicating that power over metrical frequency changed over tempo. The LMM also showed a significant main effect of *frequency* (*χ*^2^(3) = 121.90, *p* < .001).

Table S2 shows the full results of the simple effects analysis following the significant interaction. Power at the grid frequency was generally higher for slow compared to fast tempi (all *p*s < 0.038, except between neighboring tempo conditions 600 ms and 400 ms [*p* = .996] and 400 ms and 300 ms [*p* = .100]). In contrast, power at the second subharmonic was generally higher for faster than slower tempi (*p*s < .024, with the 300 ms vs 400 ms conditions contrast being the only exception [*p* = .999]). Power at the first subharmonic was lower for the 600 ms condition compared to the 300 ms, 200 ms, and 150 ms conditions (*p*s < 0.013) and for the 400 ms compared to the 200 ms condition (*p* = .003). We found no evidence that relative power at meter-unrelated frequencies differed between tempo conditions (*p*s > .509).

All meter-related frequencies displayed higher power than meter-unrelated frequencies for the 400 ms, 300 ms, and 200 ms conditions (*p*s < .002), suggesting significant synchronization at multiple metrical levels in the EEG signal. In the slowest 600 ms condition, only power at the grid frequency was higher than at unrelated frequencies (*p* < .001). In contrast, in the fastest 150 ms condition, there was significant evidence for this enhancement only for the first and second subharmonic (*p*s < .001).

Figure 3A and 3B show average *z*-scored power at the grid frequency, first and second subharmonics, and unrelated frequencies, separately for the two attention conditions. The corresponding power spectra can be found in Figure S5A and S5G. The LMM showed no evidence for a *frequency x attention* interaction, indicating that relative power at frequencies of interest did not differ significantly between attention condition (*χ*^2^(3) = 2.33, *p* = 0.507). There was also no evidence that the change of power distribution over tempo was dependent on attention (*frequency x tempo x attention* interaction; *χ*^2^(12) = 6.83, *p* = 0.869).

**Figure 3.**
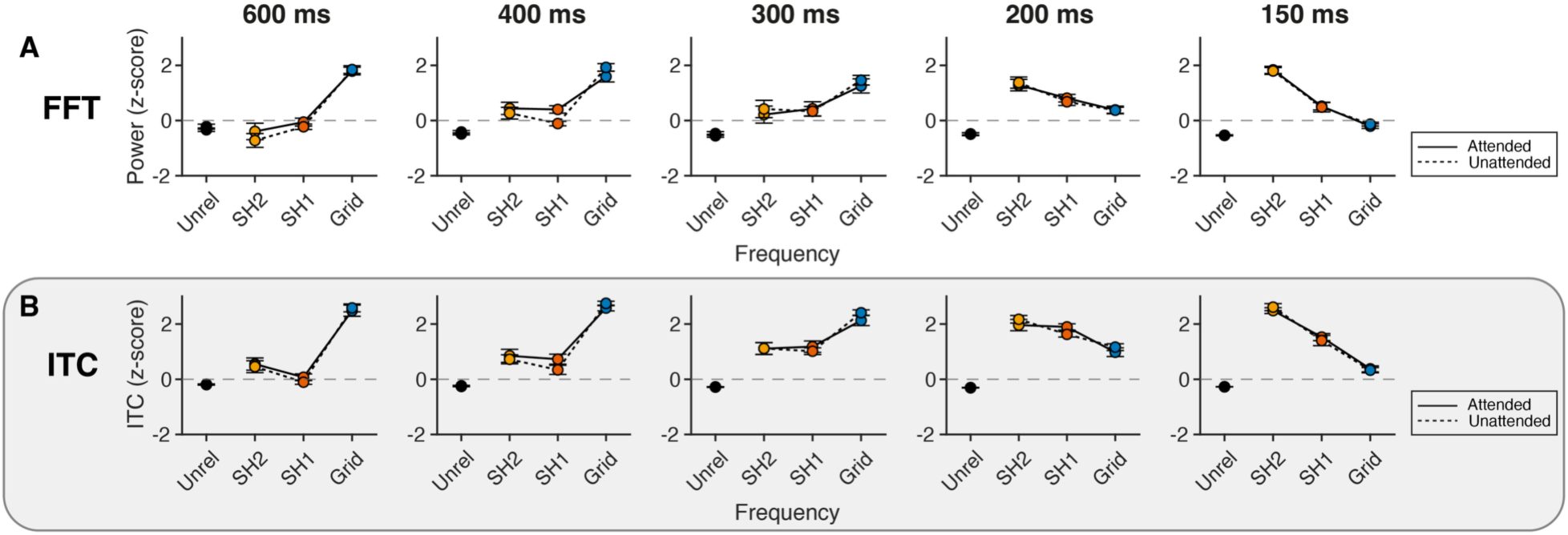
Selective enhancement of spectral power at the beat frequency was not affected by attention. (A) Average relative spectral power and (B) inter-trial phase coherence of the EEG for the attended and unattended conditions. Error bars represent standard error of the mean.

**Figure 4.**
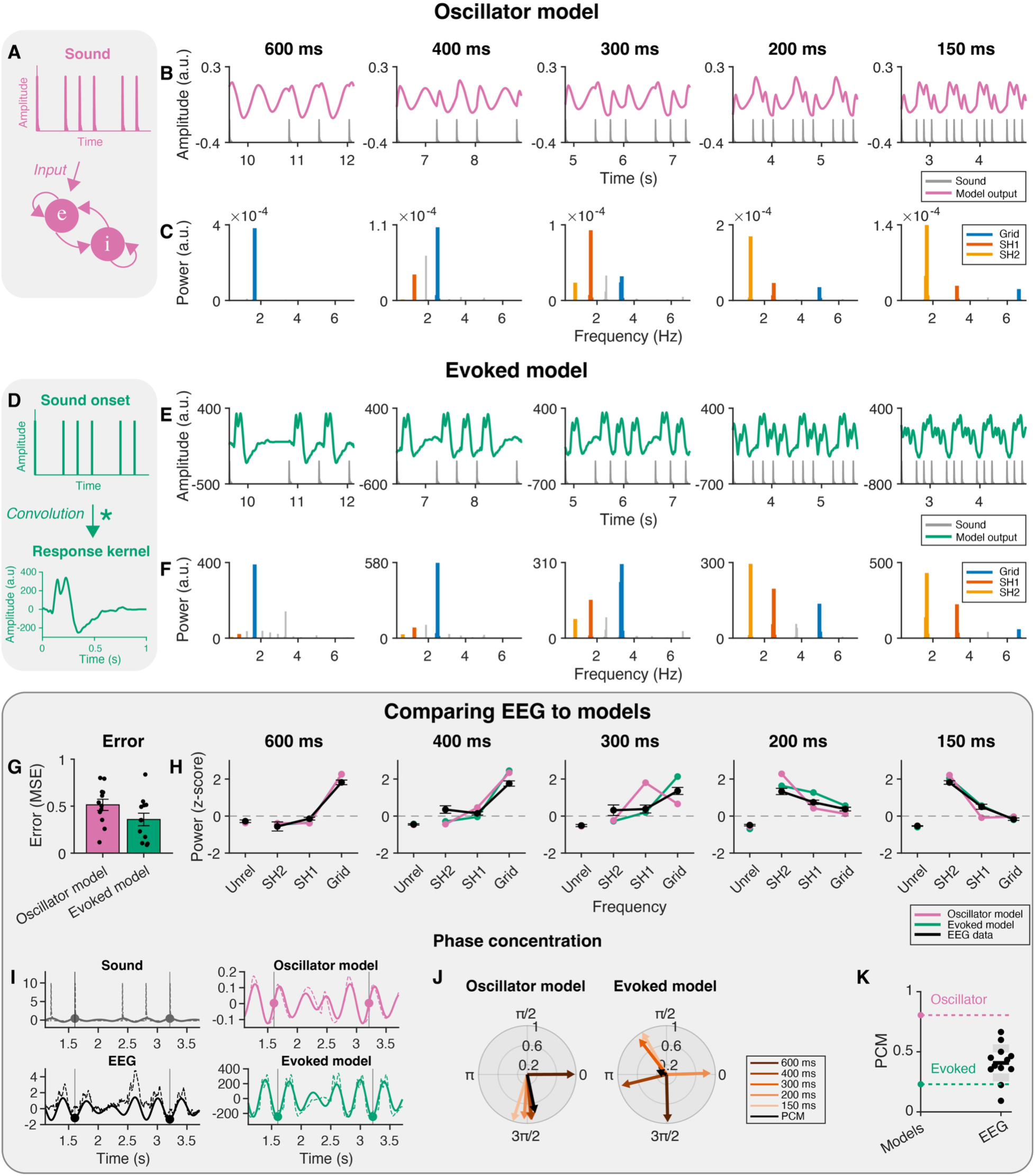
An oscillator and an evoked model capture selective enhancement of power at the beat frequency. Oscillator and evoked model description, output, and spectral power, and comparison of the models to the EEG data. (A) The oscillator model consisted of a Wilson-Cowan model responding to the continuous sound signal. (B) Example of the oscillator model output for the different tempi. Grey lines represent the sound input. Note that these examples show a 2.5 s window starting from the 5^th^ repetition of the rhythmic pattern in a block. (C) Spectral power of the oscillator model output. Note that the range of the y-axis is adapted for each tempo to depict the relative power at meter-related frequencies more clearly. (D) The evoked model consisted of a convolution of a binary sound onset signal with a response kernel. The response kernel was calculated as the EEG temporal response function over fronto-central electrodes. (E) Example of the evoked model output. Grey lines represent the sound input. (F) Spectral power of the evoked model output. (G) Mean squared error (MSE) of the oscillator and evoked model relative to the EEG data. Black dots represent individual participants and error bars represent standard error of the mean. (H) Relative (z-scored) power at meter-unrelated and meter-related frequencies for the EEG data (error bars represent standard error of the mean) and the two models. (I) Illustration of the phase concentration method for the 400 ms condition. Dashed lines represent the original sound, EEG, evoked and oscillator model signals, while the solid lines represent the signal filtered at the grid frequency (here, 2.5 Hz). The phase of the filtered signal is extracted at pattern onsets (vertical lines and corresponding markers). (J) Phase at pattern onset for all tempo conditions and phase concentration for the oscillator model and the evoked model. Angle and length of the colored vectors represent the phase and strength of synchrony, respectively. Phase concentration (PCM) is represented by the length of the black vector. (K) PCM for the individual participants (black dots) relative to the evoked model (green dashed line) and oscillator model (pink dashed line).

In a supplementary analysis, we tested whether meter-related spectral power in the EEG predicted participants’ preferred synchronization rates in the tapping task. The results showed that there was no evidence for this relationship (see Supplementary Materials section 1 for the full linear regression model results).

In summary, the FFT results show that while synchronization at the grid frequency was strongest for slow tempi, subharmonics in the EEG signal were enhanced for faster tempi. This finding is in line with our hypothesis that listeners shift the perceived beat level with changes in tempo, as was also demonstrated in the behavioral data. There was no evidence that the tempo-dependent effect differed between attention conditions.

#### ITC

Figure 2D shows the inter-trial phase coherence (ITC) per tempo condition. Figure 2E shows the *z*-scored ITC at meter-related and meter-unrelated frequencies and Figure S4B shows the *z*-scored ITC per participant. The slowest tempo (i.e., the 600 ms condition) showed strongest phase coherence at the grid frequency, while faster tempi showed more consistent phase at lower subharmonics. The LMM reflected this *tempo x frequency* interaction (*χ*^2^(12) = 434.94, *p* < .001). Additionally, there was a significant effect of *frequency* (*χ*^2^(3) = 241.23, *p* < .001).

Table S3 shows the results of the contrast analysis. ITC at the grid frequency was generally higher for slower compared to faster tempo conditions (all *p*s < .034), except between the 600 ms conditions and the 400 ms and 300 ms conditions (*p* = .882 and *p* = .298, respectively). ITC values at the first subharmonic were generally higher for faster tempo conditions (all *p*s < .001), with the exception of the differences between the 150 ms condition and the 200 ms and 300 ms conditions (*p* = .191 and *p* = .058, respectively). The second subharmonic similarly showed generally higher ITC for faster tempi (all *p*s < .001), except for the contrasts between the 400 ms condition and the 300 ms and 600 ms conditions (*p* = .119 and *p* = .239, respectively). There was no evidence for differences in ITC of unrelated frequencies between the different tempo conditions (*p*s > .912). Contrasts showed that the three meter-related frequencies in all tempo conditions were higher than meter-unrelated frequencies (*p*s < .001), except for the first subharmonic in the 600 ms condition (*p* = .584).

Figure 3A and 3B show the z-scored inter-trial phase coherence averaged over time (i.e.., over the course of the pattern), separately for attended and unattended conditions. Corresponding plots of the original ITC values unfolding over time can be found in Figure S5B and S5H. As can be observed in these figures, the ITC showed a similar pattern in the two attention conditions. The LMM for the *z*-scored ITC indeed showed no evidence for a *frequency x attention* interaction, indicating that relative phase locking at frequencies of interest was not significantly affected by attention (*χ*^2^(3) = 0.88, *p* = 0.831). Moreover, there was no evidence that the change that the ITC frequency distribution displayed over tempo differed between attention conditions (*frequency x tempo x attention* interaction: *χ*^2^(12) = 3.98, *p* = 0.984).

In summary, similar to the FFT, the ITC showed evidence for significant phase locking at meter-related frequencies, and coherence was highest at the grid frequency for slow tempo conditions and shifted to subharmonics for faster tempi. Thus, across both metrics, the EEG signal showed a pattern similar to the tapping behavior: the most prominent frequencies in the signal shifted with the tempo of the rhythmic sequences, to rates close to the preferred beat rate.

### Comparing EEG to models

#### FFT

The EEG data was tested against the oscillator model and the evoked model by comparing the change in relative power over tempi. Figure 4B and 4E show output examples of the oscillator and evoked models, respectively, and Figure 4C and 4F show their corresponding power spectra. The evoked model captured two key features of the EEG data: first, it displayed enhancement at meter-related frequencies, and second, it showed a shift over tempo from a dominant grid frequency for the slowest tempo to lower subharmonics for fast tempi (Figure 4H). The oscillator model also fits the overall pattern of the data, similarly showing selectively enhanced subharmonics relative to the grid frequency for faster but not slower tempi (Figure 4H). Quantifying these fits, Figure 4G shows the mean squared error (MSE) of both models relative to the EEG data. A Bayesian *t*-test showed anecdotal evidence for a larger error for the oscillator model compared to the evoked model (BF_10_ = 1.42, *t*(11) = 2.09, *p* = .061) with a moderate effect size (*d* = 0.60). Hence, there is no strong preference for one model over the other based on these data.

#### Phase concentration

Figure 4J shows the average phase lag between the output signals of the oscillator and evoked models and the sound signal for every tempo condition separately, with the overall phase concentration metric (PCM) represented by the length of the black arrow. As expected, the oscillator model showed higher phase concentration (*PCM_O_* = 0.80) than the evoked model (*PCM_E_* = 0.23). Similarly, the PCM was quantified for the EEG signal of every participant (see Figure S6 for individual polar plots). The individual PCM values (*PCM_EEG_* mean = 0.41, SD = 0.15) are shown in Figure 4K (black points). We compared the squared error of these values relative to the PCM of each of the models using a paired samples Bayesian *t*-test, which showed that the MSE for the oscillator model was larger than the MSE for the evoked model (i.e., the PCM of the EEG data was closer to the evoked model than to the oscillator model; BF_10_ = 2.47, *t*(11) = 2.49, *p* = .030, with a moderate effect size of *d* = 0.72). However, additional Bayesian *t*-tests showed that the PCM values of the EEG fell in between the two models: they were higher than expected from the evoked model (BF_10_ = 21.88, *t*(11) = 4.01, *p* = .002, with a large effect size of *d* = 1.16), while there was also decisive evidence that they were lower than the oscillator model (BF_10_ > 100, *t*(11) = −8.99, *p* < .001, with a large effect size of *d* = −2.60).

To test whether attention affected the PCM, we performed Bayesian *t*-tests comparing the PCM of the participants’ EEG against the two models separately for the attended and unattended conditions (Figure S5F and S5L). These showed that the EEG’s PCM was higher than the evoked model in both the attended (BF_10_ > 100, *t*(11) = 5.14, *p* < .001, *d* = 1.48) and the unattended conditions (BF_10_ = 2.29, *t*(11) = 2.44) *p* = .033, *d* = 0.70). There was also decisive evidence that the participants’ PCM were lower than the oscillator model in both attention conditions (BFs_10_ > 100, *t*s(11) < −7.19, *p*s < .001, *d*s < 2.07). Overall, these findings indicate that the phase concentration of the EEG signal fell between the PCM predicted by the two model, irrespective of active attention.

## Discussion

The current study investigated whether neural responses to rhythm show evidence of entrainment by testing how EEG responses change over tempo. We showed that neural synchronization to rhythm did not scale with tempo, but was instead selectively enhanced at the level of the beat (i.e., around 2 Hz), regardless of whether the rhythm was actively attended. Neural spectral power and phase locking mirrored the tapping results, in which participants switched to a slower metrical level for faster tempi. Our findings suggest a preferential frequency range of neural synchronization in the auditory-motor system, which coincides with the preferred 1-2 Hz rate of beat perception, spontaneous motor behavior, and optimal attentional sampling (Moelants 2002; MacDougall and Moore 2005; McAuley et al. 2006; Zalta et al. 2020; Weineck et al. 2022).

These results may indicate that beat perception is supported by an oscillatory mechanism, since the tempo-dependent selective enhancement at the beat frequency is not present in the sound itself: it is in the transformation from the sound to cortical neural signals that the internal beat is represented (Nozaradan, Keller, et al. 2018; Lenc et al. 2021). In this way, a pattern played at different tempi may give rise to a similar beat percept, reflecting a many-to-one mapping between input and neural representation (Nozaradan, Keller, et al. 2018). In line with this hypothesis, we found that the consistent enhancement of beat-related frequencies in the EEG signal could be mimicked by an oscillator model.

However, this selective enhancement could also be replicated by a model of passive evoked neural responses to the individual sounds. We show that lower metrical frequencies can emerge from the way evoked responses interact when they track a rhythmic sequence, even in the absence of genuine entrainment. This may be due to the overlap between the frequency content of slower auditory evoked responses and the delta range (0.5-4 Hz) (Başar-Eroglu et al. 1992; Güntekin and Başar 2016), at which a beat is usually perceived. The convolution of a rhythmic input with an evoked response acts like a filter, enhancing those harmonics of the input that fall within the dominant frequency range of the evoked response in the delta band. In this way, subharmonics that are not strongly represented in the sound input can nonetheless become dominant in the neural response, indicating that selective enhancements in the frequency-domain cannot straightforwardly be interpreted as oscillatory activity in the narrow sense (Capilla et al. 2011; Novembre and Iannetti 2018; Oganian et al. 2023). These findings highlight that the dynamics of evoked responses should be considered when interpreting neural entrainment to rhythmic stimuli by, for example, explicitly using an evoked model as a baseline for explaining neural synchronization, or alternatively, relying on identical stimuli with different subjectively induced meters (e.g., Nozaradan et al. 2011; Nave et al. 2022).

As expected, we found that the oscillator and evoked models made different predictions about phase concentration: because the evoked response always occurred at the same time lag after sound onset, the predicted phase concentration over different tempi was low for the evoked model output, while it was relatively high for the oscillator model output. The phase concentration of the EEG data was closest to the evoked model. This is in contrast to Doelling et al. (2019), who found that MEG responses to isochronous rhythms were best supported by an oscillator model. This discrepancy was predicted by the preceding work, which showed that brain dynamics are more oscillatory when acoustic edges (e.g., note onsets) are smoother, suggesting that the sharper onsets in our experiment would in turn yield stronger evoked dynamics. Our findings support this prediction, suggesting oscillatory activity may be enhanced in more realistic musical pieces. While the neural phase concentration was closest to the evoked model, it still fell in between the two model predictions: it was larger than expected from the evoked model alone, but smaller than predicted by the oscillator model. While we did not explicitly test this, it is possible that the neural signal contains contributions from both transient responses and oscillatory entrainment, as suggested previously (Doelling et al. 2019). Future research could aim to quantify these separate contributions.

Compared to Doelling et al. (2019), there were some differences in how the evoked model predictions were generated. In Doelling et al. (2019), the evoked responses were measured separately at the start of the experiment which may have been less representative for the neural responses throughout the longer musical excerpt. Here, we extracted this evoked response from data during the rhythm presentation, incorporating responses at different tempi and stages within the block, thereby capturing potential effects of habituation and refraction. Whether this difference significantly affects the prediction of the evoked model is debatable; the two models in both papers yield similar predictions for a similar range of stimulus rates. Our evoked response method also increases the risk of capturing ongoing oscillatory activity in the response kernel, which could in turn drive the enhancement of metrical frequencies in the evoked model. However, a model in which the evoked response was based on meter-unrelated tempo conditions (that is, the 150, 300, and 600 ms conditions were used to create an evoked model for the 200 and 400 ms conditions, and vice versa) showed the same patterns of results, confirming that the response kernel captured a generalized response to the sounds.

The goal of the current study was to distinguish between entrainment and evoked responses based on two pre-defined models that are simple instances of their class (after Doelling et al. 2019). However, to arrive at a best-fitting model of the neural data, future studies could consider using more complex models. For example, a more realistic evoked model could incorporate additional features that modulate the amplitude of ERP components, such as the effect of inter-onset-interval, as well as habituation over longer sequences (Pereira et al. 2014; Damsma et al. 2021). These parameters could be based on an independent dataset prior to computing an evoked model. Similarly, more complex non-linear models of entrainment, such as gradient frequency neural networks, could be considered, as they have been shown to capture beat and meter perception (Large et al. 2015; Large et al. 2023).

The modeling results show that evoked responses alone result in selective enhancement at the beat frequency. This does not rule out, however, that oscillatory and evoked responses may arise from the same neural circuitry and, therefore, interact. While we modeled oscillatory and evoked responses independently, cyclical changes in neuronal excitability have been hypothesized to amplify or suppress stimulus processing (Schroeder and Lakatos 2009), and ongoing low-frequency oscillations indeed modulate event-related cortical responses (Lakatos et al. 2005; Lakatos et al. 2007). Conversely, our results also raise the possibility that evoked responses themselves play a role in the interpretation of beat and meter by means of ‘neural emphasis’ at specific positions in a rhythmic sequence, and could drive attentional sensitivity through bottom-up fluctuations in cortical excitability (Rajendran et al. 2017; Rajendran et al. 2020). Future studies could test these interactions by investigating the relationship between neural response magnitude and perceived beat location.

The enhancement of low-frequency subharmonics could alternatively be explained by the commonly observed decrease in power of neural activity as a function of frequency, known as 1/*f* activity (Pritchard 1992). However, this general profile is unlikely to cause the current findings. First, the analyses focused on phase-locked activity by removing 1/*f* background activity through the subtraction of neighboring values from each frequency point (e.g., Nozaradan et al. 2011) and by analyzing phase consistency (ITC). While 1/*f* background activity could affect ITC values, more noise at low frequencies would cause reduced ITC in the low end and enhanced ITC in the high end: the opposite of what we find in the fast tempo conditions. Second, the results show selective enhancement at the preferred beat rate. If the enhancement would be driven by the 1/*f* profile, one would expect the lowest frequencies to always be strongest. Finally, the selective enhancement could be mimicked by an evoked model. The model was created by convolving a response kernel with a time series representing the sound onsets and should therefore not be subject to the power-law profile of EEG background activity.

The results showed no difference in neural responses between conditions where participants were instructed to actively attend the rhythm and passive listening conditions. This suggests that the observed transformation between the power spectrum of the sound input and the neural response does not require top-down attention. Regardless of whether neural responses are attributed to evoked or oscillatory activity, these results indicate that the cortex passively tracks rhythmic sequences, in line with previous studies showing that task relevance is not necessary in forming temporal expectations (Ladinig et al. 2009; Bouwer et al. 2014; Damsma and van Rijn 2017; Bouwer et al. 2020). Several recent papers did find persistent oscillations after a rhythmic stimulus was stopped (Kösem et al. 2018; van Bree et al. 2021; Bouwer et al. 2023), which is sometimes seen as evidence for entrainment (although see Doelling and Assaneo 2021), but in all these cases, the rhythm was task-relevant and attended to. Without task-relevance, such sustained entrainment was not observed (Lerousseau et al. 2021). This may further suggest that the cortical selective enhancement we observed, which was independent of attention, was caused by overlapping evoked responses, rather than entrainment. However, recent work suggests that attention may still enhance steady-state evoked responses, but that this is limited to certain types of rhythms (i.e., rhythms with a strong beat) and entraining frequencies (i.e., higher beat-related frequencies) (Gibbings et al. 2023).

In addition, Gibbings et al. (2023) found that steady-state evoked responses did not predict individuals’ tapping rates, which is in line with our current results. While our results exhibited substantial variation in dominant tapping rates between individuals, these did not correlate with neural synchronization at meter-related frequencies. These findings raise the question of whether the frequency at which neural entrainment is strongest determines the subjective experience of beat and meter. The direct relation between frequency tagging and beat perception has been debated (Henry et al. 2017; Rajendran and Schnupp 2019). On the one hand, individual differences in the amplitude of neural entrainment at beat frequencies have been linked to the accuracy of sensorimotor synchronization and temporal predictions of upcoming events (Nozaradan et al. 2016). This was specifically the case for slower tempi (under 2 Hz), potentially because attention switches to lower metrical levels for fast tempi, as we demonstrated in the current study. In addition, the phase of neural delta entrainment has been shown to predict auditory sensitivity (Kaya and Henry 2022). On the other hand, subjective ratings of beat strength have been shown to be independent from neural power at beat-related frequencies relative to the stimulus (Henry et al. 2017). Future studies with a large sample size are needed to further test the relationship between individual differences in preferred tapping rate and neural synchronization. The goal of our study design was to increase power and decrease bias by testing a very high number of trials per participant and condition (i.e., more than 4500 pattern repetitions over the course of the experiment) (Rouder et al. 2023). This approach aimed to avoid replicability issues by precisely characterizing the performance of each participant separately, while a large sample size does not always lead to more accurate results and could even breed confidence in a wrong answer in conditions with high trial noise (Meng 2018; Rouder et al. 2023). However, attentional entrainment in a behavioral paradigm has been shown to vary widely between individuals (Sun et al. 2022). Thus, even though the shift in neural synchronization over tempi was relatively consistent in the current sample, a larger sample size is necessary to characterize between-person differences and generalize our findings.

In conclusion, we showed that the selective enhancement of cortical responses to rhythm at the beat frequency is tempo-dependent, similar to the behavioral shift in the hierarchical level listeners tap to when responding to rhythm. This effect was independent of attention. While cortical responses to rhythm thus mirror the perceived beat, and not necessarily the sound input, we show that these results can be mimicked by both an oscillatory model and a model of evoked responses. Thus, while the selective enhancement of cortical responses at the beat frequency may be interpreted as oscillatory entrainment, it could also result from the succession of evoked responses.

## Data availability

All data and code used for data acquisition and analysis are available at https://osf.io/ghp94/.

## Author contributions

**Atser Damsma:** Conceptualization, Data curation, Formal analysis, Methodology, Project administration, Visualization, Writing – original draft, Writing – review & editing. **Mitchell de Roo:** Formal analysis, Methodology, Writing – review & editing. **Keith Doelling:** Methodology, Software, Writing – review & editing. **Pierre-Louis Bazin:** Funding acquisition, Methodology, Writing – review & editing. **Fleur L. Bouwer:** Conceptualization, Data curation, Formal analysis, Funding acquisition, Investigation, Methodology, Project administration, Supervision, Visualization, Writing – review & editing.

