## Supplementary Materials for "Tempo-dependent selective enhancement of neural responses at the beat frequency can be mimicked by both an oscillator and an evoked model"

### 1. EEG and preferred tapping rate

If oscillatory neural entrainment underlies beat perception, we might expect a relationship between participants' meter-related EEG activity and their preferred tapping rate. To test this hypothesis, we tested whether relative, *z*-scored power at meter-related frequencies (i.e., grid frequency, first sub-harmonic, and second sub-harmonic) of the EEG signal (Figure S4A) predicted relative power at these frequencies in the tapping signal (Figure S3B). A linear regression model was computed with *relative tapping power* as the dependent variable and *frequency*, *tempo*, their interaction and *relative EEG power* (averaged over attention conditions) as predictors. Omnibus effects of this model were estimated with a Type III ANOVA using the *Anova* function from the *car* package in *R* (Fox et al., 2012). While the results showed significant effects of *frequency* ($F[2,161] = 4.76$, $p = .009$), *tempo* ($F[4,161] = 4.89$, $p < .001$), and their interaction ($F[8,161] = 6.63$, $p < .001$), we found no evidence for an additional predictive effect of EEG power ($F[1,161] = 0.05$, p = .830). These results suggest that the strength of neural synchronization at a specific frequency did not entail a similarly strong behavioral synchronization at that rate.

### 2. Evoked model based on frequency-unrelated tempi

To test whether the power spectrum of the evoked model depended on tempo-specific oscillations implicitly captured in the response kernel, an alternative evoked model was calculated based on two separate, tempo-unrelated response kernels. To this end, a response kernel was estimated based on EEG data for the 150, 300 and 600 ms tempo conditions and, separately, for the 200 and 400 ms conditions, using the same methods as for the original response kernel. The responses kernel computed for the 150, 300, and 600 ms tempo conditions was then used to compute an evoked model of the unrelated 200 and 400 ms conditions, and vice versa.

Figure S1 shows the two separate response kernels calculated over the 150, 300, and 600 ms tempo conditions and the 200 and 400 ms conditions, and Figure S2 shows the power spectrum of the original evoked model (Figure S2A and S2B) and the model based on these tempo-unrelated response kernels (Figure S2C and S2D). Visual inspection suggests that the results were highly similar. Indeed, like the original evoked model, a Bayesian *t*-test showed anecdotal evidence for a larger error for the oscillator model than for the evoked model ($BF_{10} = 1.27$, $t(11) = 2.00$, $p = .071$) with a moderate effect size ($d = 0.58$). The PCM of the evoked model based on frequency-unrelated tempi ($PCM_E = 0.20$) was also similar to the original model ($PCM_E = 0.21$). The PCM values of the EEG data were marginally closer to the evoked model than to the oscillator model ($BF_{10} = 1.58$, $t(11) = 2.16$, $p = .053$, $d = 0.63$), but it was also higher than expected from the evoked data alone ($BF_{10} = 55.10$, $t(11) = 4.67$, $p < .001$, $d = 1.35$). These results are qualitatively similar to the original evoked model based on all tempi, showing that the beat-specific enhancements in the power spectrum of this model are unlikely to result from oscillations captured in the response kernel.

### Supplementary Figures

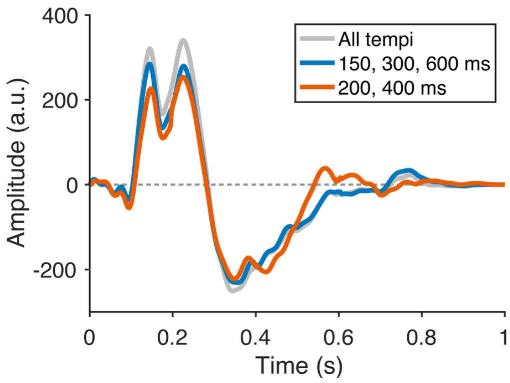


***Figure S1.*** *The original response kernel calculated over all tempi (grey line), and two separate response kernels computed over harmonically related tempi.*

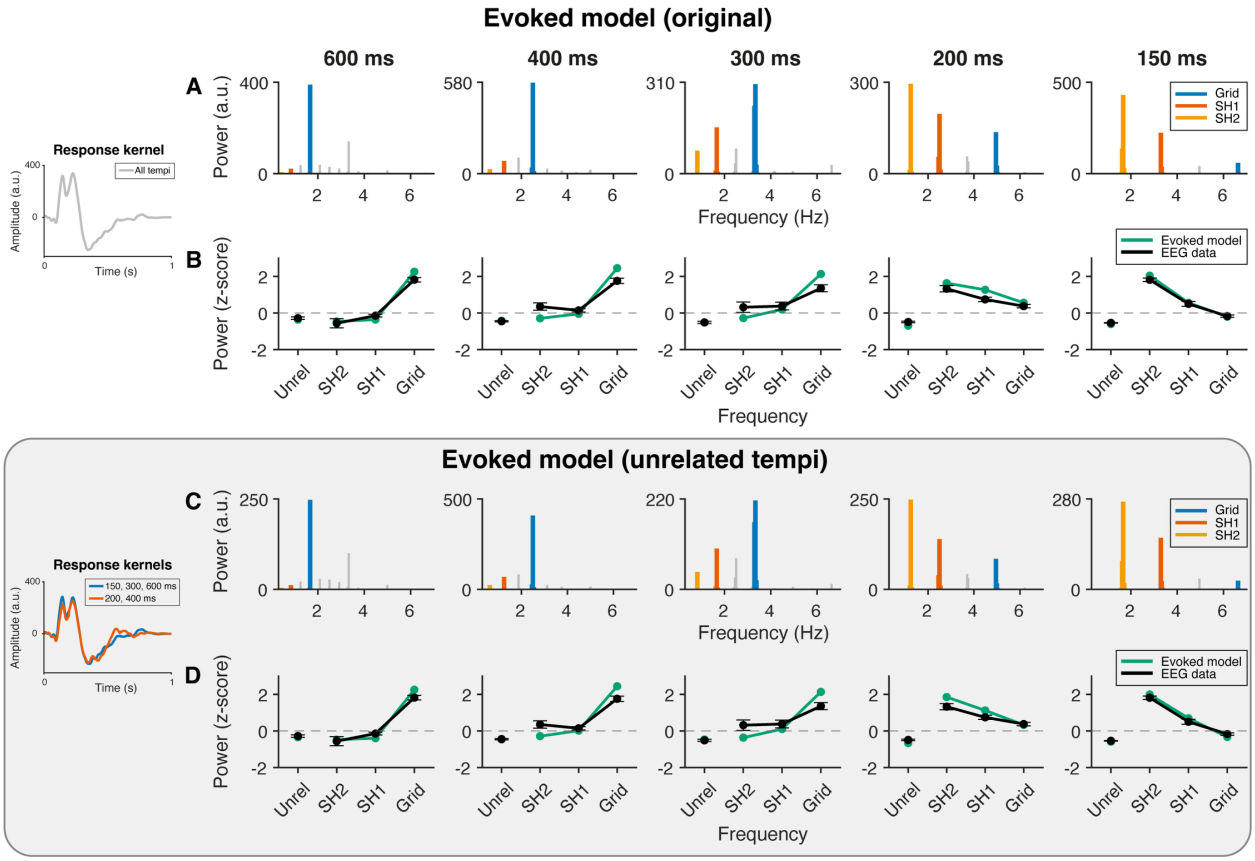


***Figure S2. Absolute and relative spectral power of the original evoked model and an alternative based on two separate response kernels, which were calculated over frequency-unrelated tempi.*** *A) Spectral power of the original evoked model. (B) Relative (z-scored) power at meter-unrelated and meter-related frequencies for the EEG data (error bars represent standard error of the mean) and the original evoked model. C) Spectral power of the evoked model based on frequency-unrelated tempi. (D) Relative power for the EEG data and the evoked model based on frequency-unrelated tempi.*

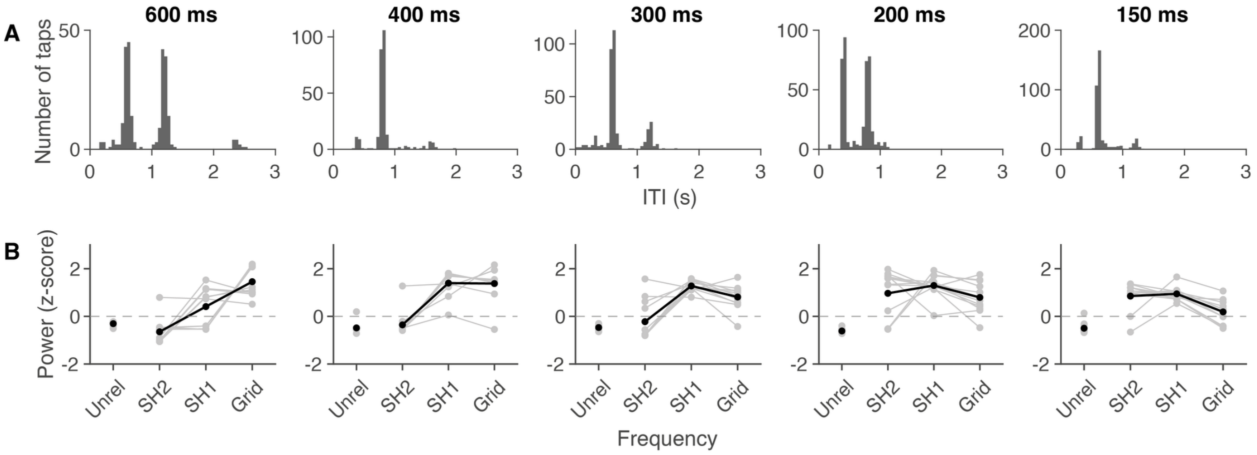


***Figure S3.*** *Tapping results. A) Histogram of the inter-tap intervals (ITI) of all participants for each tempo B) Relative (z-scored) spectral power of the tapping data for meter-unrelated frequencies, the second and first subharmonics, and the grid frequency per participant (grey lines). Black lines represent the average over participant.*

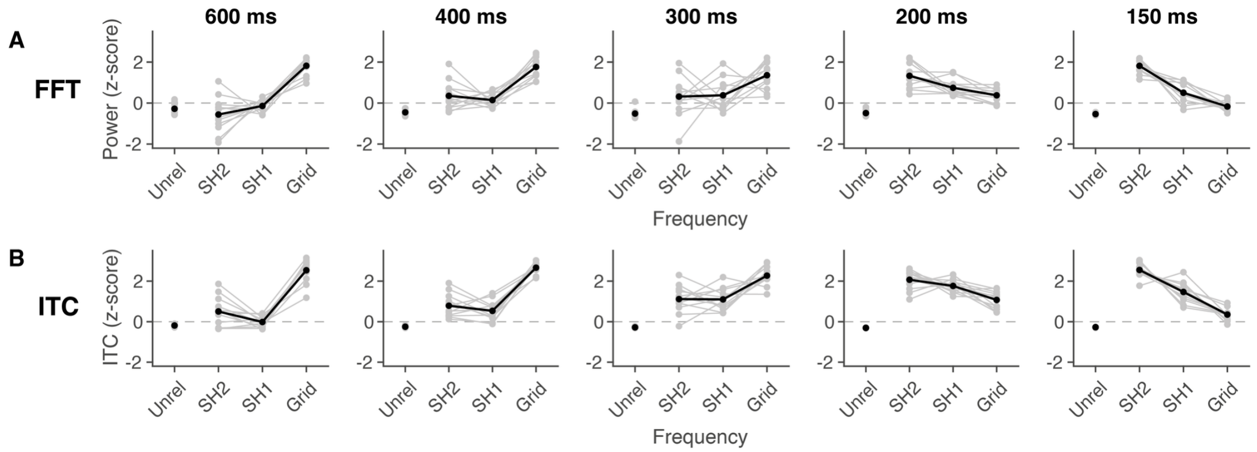


***Figure S4.*** *Individual differences in neural synchronization. (A) Relative (z-scored) spectral power (FFT) and (B) phase coherence (ITC) for meter-unrelated frequencies, the second and first subharmonics, and the grid frequency per participant (grey lines). Black lines represent the average over participants. Note that the relative power and ITC values were averaged over attention conditions.*

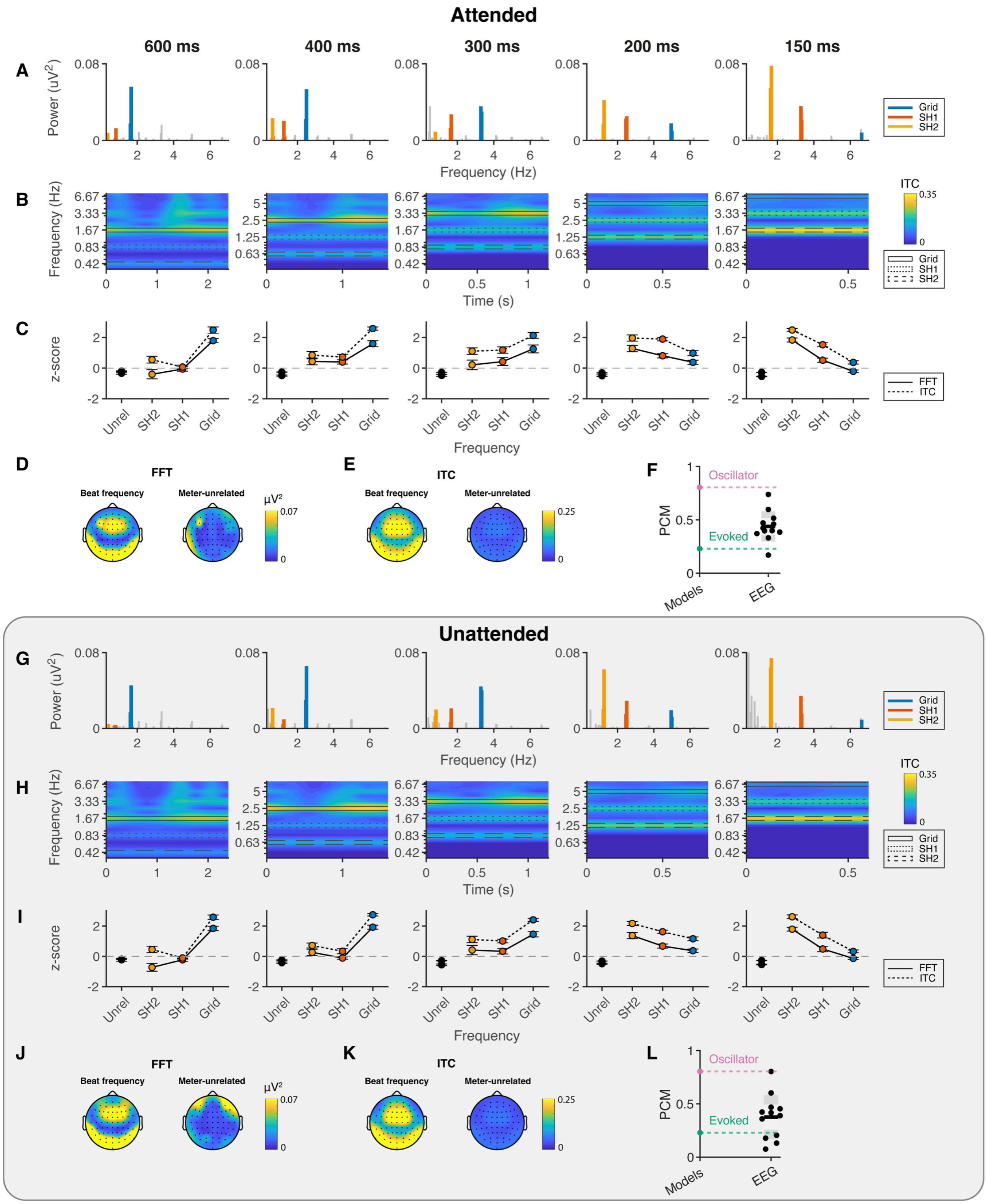


***Figure S5. Effect of attention.*** *Spectral power (A,G), inter-trial phase coherence (B,H), relative spectral power and ITC at meter-unrelated and meter-related frequencies (C,I), topographies of spectral power (D,J) and ITC (E,K) at the beat frequency and meter-unrelated frequencies for the EEG, and phase concentration (F,L), for the attended and unattended conditions, respectively.*

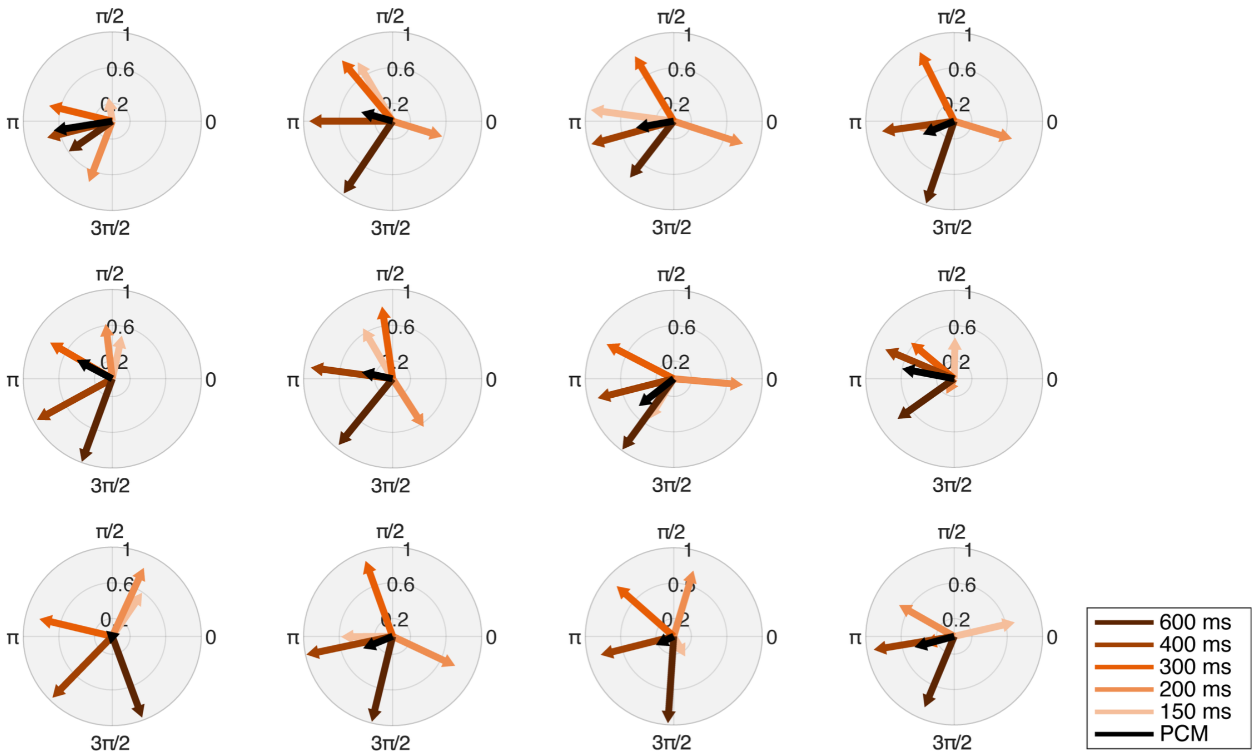


***Figure S6. Phase at pattern onset for all tempo conditions and phase concentration (PCM) for each individual participant.*** *Angle and length of the colored vectors represent the phase and strength of synchrony, respectively. Phase concentration (PCM) is represented by the length of the black vector.*

### Supplementary Tables

***Table S1. Results of the contrast analysis on z-scored spectral power of the tapping data.*** *A Tukey's pairwise multiple comparison test was calculated with corresponding standardized effect sizes (d).*

| Frequency | Contrast | Estimate | *p* | *d* |
|---|---|---|---|---|
| Unrelated frequencies | 600 ms - 400 ms | 0.18 | 0.933 | 0.33 |
| | 600 ms - 300 ms | 0.16 | 0.948 | 0.30 |
| | 600 ms - 200 ms | 0.31 | 0.636 | 0.57 |
| | 600 ms - 150 ms | 0.19 | 0.909 | 0.35 |
| | 400 ms - 300 ms | -0.02 | 1.000 | -0.03 |
| | 400 ms - 200 ms | 0.13 | 0.980 | 0.24 |
| | 400 ms - 150 ms | 0.01 | 1.000 | 0.02 |
| | 300 ms - 200 ms | 0.14 | 0.966 | 0.27 |
| | 300 ms - 150 ms | 0.03 | 1.000 | 0.05 |
| | 200 ms - 150 ms | -0.12 | 0.985 | -0.21 |
| Second subharmonic | 600 ms - 400 ms | -0.29 | 0.701 | -0.54 |
| | 600 ms - 300 ms | -0.43 | 0.310 | -0.78 |
| | 600 ms - 200 ms | -1.62 | <0.001 | -2.98 |
| | 600 ms - 150 ms | -1.50 | <0.001 | -2.77 |
| | 400 ms - 300 ms | -0.13 | 0.976 | -0.25 |
| | 400 ms - 200 ms | -1.32 | <0.001 | -2.44 |
| | 400 ms - 150 ms | -1.21 | <0.001 | -2.23 |
| | 300 ms - 200 ms | -1.19 | <0.001 | -2.19 |
| | 300 ms - 150 ms | -1.07 | <0.001 | -1.98 |
| | 200 ms - 150 ms | 0.12 | 0.985 | 0.21 |
| First subharmonic | 600 ms - 400 ms | -0.99 | <0.001 | -1.82 |
| | 600 ms - 300 ms | -0.87 | 0.001 | -1.60 |
| | 600 ms - 200 ms | -0.89 | 0.001 | -1.63 |
| | 600 ms - 150 ms | -0.54 | 0.115 | -0.99 |
| | 400 ms - 300 ms | 0.12 | 0.985 | 0.22 |
| | 400 ms - 200 ms | 0.10 | 0.992 | 0.18 |
| | 400 ms - 150 ms | 0.45 | 0.275 | 0.83 |
| | 300 ms - 200 ms | -0.02 | 1.000 | -0.04 |
| | 300 ms - 150 ms | 0.33 | 0.566 | 0.61 |
| | 200 ms - 150 ms | 0.35 | 0.510 | 0.65 |
| Grid frequency | 600 ms - 400 ms | 0.08 | 0.997 | 0.14 |
| | 600 ms - 300 ms | 0.64 | 0.033 | 1.18 |
| | 600 ms - 200 ms | 0.66 | 0.027 | 1.22 |
| | 600 ms - 150 ms | 1.27 | <0.001 | 2.34 |
| | 400 ms - 300 ms | 0.57 | 0.096 | 1.04 |
| | 400 ms - 200 ms | 0.58 | 0.080 | 1.07 |
| | 400 ms - 150 ms | 1.19 | <0.001 | 2.19 |
| | 300 ms - 200 ms | 0.02 | 1.000 | 0.03 |
| | 300 ms - 150 ms | 0.63 | 0.041 | 1.15 |
| | 200 ms - 150 ms | 0.61 | 0.051 | 1.12 |

***Table S2. Results of the contrast analysis on z-scored spectral power (FFT) of the EEG data.*** *A Tukey's pairwise multiple comparison test was calculated with corresponding standardized effect sizes (d).*

| Frequency | Contrast | Estimate | *p* | *d* |
|---|---|---|---|---|
| Unrelated frequencies | 600 ms - 400 ms | 0.17 | 0.828 | 0.31 |
| | 600 ms - 300 ms | 0.23 | 0.605 | 0.41 |
| | 600 ms - 200 ms | 0.21 | 0.696 | 0.37 |
| | 600 ms - 150 ms | 0.26 | 0.509 | 0.46 |
| | 400 ms - 300 ms | 0.06 | 0.996 | 0.11 |
| | 400 ms - 200 ms | 0.04 | 0.999 | 0.07 |
| | 400 ms - 150 ms | 0.09 | 0.985 | 0.15 |
| | 300 ms - 200 ms | -0.02 | 1.000 | -0.04 |
| | 300 ms - 150 ms | 0.02 | 1.000 | 0.04 |
| | 200 ms - 150 ms | 0.05 | 0.998 | 0.08 |
| Second subharmonic | 600 ms - 400 ms | -0.91 | <0.001 | -1.62 |
| | 600 ms - 300 ms | -0.88 | <0.001 | -1.55 |
| | 600 ms - 200 ms | -1.89 | <0.001 | -3.35 |
| | 600 ms - 150 ms | -2.38 | <0.001 | -4.21 |
| | 400 ms - 300 ms | 0.04 | 0.999 | 0.06 |
| | 400 ms - 200 ms | -0.98 | <0.001 | -1.73 |
| | 400 ms - 150 ms | -1.46 | <0.001 | -2.59 |
| | 300 ms - 200 ms | -1.01 | <0.001 | -1.79 |
| | 300 ms - 150 ms | -1.50 | <0.001 | -2.66 |
| | 200 ms - 150 ms | -0.49 | 0.024 | -0.86 |
| First subharmonic | 600 ms - 400 ms | -0.29 | 0.394 | -0.51 |
| | 600 ms - 300 ms | -0.52 | 0.013 | -0.92 |
| | 600 ms - 200 ms | -0.89 | <0.001 | -1.57 |
| | 600 ms - 150 ms | -0.64 | 0.001 | -1.14 |
| | 400 ms - 300 ms | -0.23 | 0.607 | -0.41 |
| | 400 ms - 200 ms | -0.60 | 0.003 | -1.06 |
| | 400 ms - 150 ms | -0.36 | 0.187 | -0.63 |
| | 300 ms - 200 ms | -0.36 | 0.168 | -0.65 |
| | 300 ms - 150 ms | -0.12 | 0.944 | -0.22 |
| | 200 ms - 150 ms | 0.24 | 0.573 | 0.43 |
| Grid frequency | 600 ms - 400 ms | 0.06 | 0.996 | 0.11 |
| | 600 ms - 300 ms | 0.46 | 0.038 | 0.82 |
| | 600 ms - 200 ms | 1.44 | <0.001 | 2.55 |
| | 600 ms - 150 ms | 1.99 | <0.001 | 3.52 |
| | 400 ms - 300 ms | 0.40 | 0.100 | 0.71 |
| | 400 ms - 200 ms | 1.38 | <0.001 | 2.45 |
| | 400 ms - 150 ms | 1.93 | <0.001 | 3.42 |
| | 300 ms - 200 ms | 0.98 | <0.001 | 1.73 |
| | 300 ms - 150 ms | 1.53 | <0.001 | 2.70 |
| | 200 ms - 150 ms | 0.55 | 0.008 | 0.97 |

***Table S3. Results of the contrast analysis on z-scored inter-trial phase coherence (ITC) of the EEG data.*** *A Tukey's pairwise multiple comparison test was calculated with corresponding standardized effect sizes (d).*

| Frequency | Contrast | Estimate | *p* | *d* |
|---|---|---|---|---|
| Unrelated frequencies | 600 ms - 400 ms | 0.06 | 0.992 | 0.13 |
| | 600 ms - 300 ms | 0.09 | 0.963 | 0.19 |
| | 600 ms - 200 ms | 0.12 | 0.912 | 0.25 |
| | 600 ms - 150 ms | 0.08 | 0.973 | 0.18 |
| | 400 ms - 300 ms | 0.03 | 0.999 | 0.07 |
| | 400 ms - 200 ms | 0.06 | 0.993 | 0.12 |
| | 400 ms - 150 ms | 0.02 | 1.000 | 0.05 |
| | 300 ms - 200 ms | 0.03 | 1.000 | 0.06 |
| | 300 ms - 150 ms | -0.01 | 1.000 | -0.01 |
| | 200 ms - 150 ms | -0.03 | 0.999 | -0.07 |
| Second subharmonic | 600 ms - 400 ms | -0.28 | 0.239 | -0.60 |
| | 600 ms - 300 ms | -0.61 | <0.001 | -1.29 |
| | 600 ms - 200 ms | -1.56 | <0.001 | -3.31 |
| | 600 ms - 150 ms | -2.05 | <0.001 | -4.33 |
| | 400 ms - 300 ms | -0.33 | 0.119 | -0.69 |
| | 400 ms - 200 ms | -1.28 | <0.001 | -2.71 |
| | 400 ms - 150 ms | -1.77 | <0.001 | -3.73 |
| | 300 ms - 200 ms | -0.96 | <0.001 | -2.02 |
| | 300 ms - 150 ms | -1.44 | <0.001 | -3.04 |
| | 200 ms - 150 ms | -0.48 | 0.004 | -1.02 |
| First subharmonic | 600 ms - 400 ms | -0.55 | 0.001 | -1.16 |
| | 600 ms - 300 ms | -1.11 | <0.001 | -2.35 |
| | 600 ms - 200 ms | -1.78 | <0.001 | -3.75 |
| | 600 ms - 150 ms | -1.48 | <0.001 | -3.13 |
| | 400 ms - 300 ms | -0.57 | <0.001 | -1.19 |
| | 400 ms - 200 ms | -1.23 | <0.001 | -2.60 |
| | 400 ms - 150 ms | -0.93 | <0.001 | -1.97 |
| | 300 ms - 200 ms | -0.66 | <0.001 | -1.40 |
| | 300 ms - 150 ms | -0.37 | 0.058 | -0.77 |
| | 200 ms - 150 ms | 0.30 | 0.191 | 0.63 |
| Grid frequency | 600 ms - 400 ms | -0.13 | 0.882 | -0.27 |
| | 600 ms - 300 ms | 0.27 | 0.298 | 0.56 |
| | 600 ms - 200 ms | 1.47 | <0.001 | 3.10 |
| | 600 ms - 150 ms | 2.18 | <0.001 | 4.61 |
| | 400 ms - 300 ms | 0.39 | 0.034 | 0.83 |
| | 400 ms - 200 ms | 1.59 | <0.001 | 3.37 |
| | 400 ms - 150 ms | 2.31 | <0.001 | 4.88 |
| | 300 ms - 200 ms | 1.20 | <0.001 | 2.54 |
| | 300 ms - 150 ms | 1.92 | <0.001 | 4.05 |
| | 200 ms - 150 ms | 0.72 | <0.001 | 1.52 |
